# LONG-TERM MONITORING OF THE EUROPEAN ROLLER (*CORACIAS GARRULUS*) IN UKRAINE: IS CLIMATE BEHIND THE CHANGES?

**DOI:** 10.1101/2021.12.17.473117

**Authors:** T. Shupova, V. Tytar

## Abstract

Since the 1980s there has been a long-term decline in numbers and contraction of range in Europe, including Ukraine. Our specific goals were to reconstruct the climatically suitable range of the species in Ukraine before the 1980s, gain better knowledge on its requirements, compare the past and current suitable areas, infer the regional and environmental variables that best explain its occurrence, and quantify the overall range change in the country. For these purposes we created a database consisting of 347 records of the roller made ever in Ukraine. We employed a species distribution modeling (SDM) approach to hindcast changes in the suitable range of the roller during historical times across Ukraine and to derive spatially explicit predictions of climatic suitability for the species under current climate. SDMs were created for three time intervals (before 1980, 1985-2009, 2010-2021) using corresponding climate data extracted from the TerraClim database. SDMs show a decline of suitable for rollers areas in the country from 85 to 46%. Several factors, including land cover and use, human population density and climate, that could have contributed to the decline of the species in Ukraine were considered. We suggest climate change and its speed (velocity) have been responsible for shaping the contemporary home range of the European roller.

## Introduction

The European roller (*Coracias garrulus*) is the only member of the roller family of birds to breed in Europe. Being a bird of warmer regions, its overall range extends into the Middle East, Central Asia and Morocco. The species is commonly met in dry, open country with scattered trees, preferring lowlands (Cramp 1985). The European roller is a long-distance migrant, wintering in Africa south of the Sahara. In Ukraine, they arrive at nesting sites in late April - early May. The bird, particularly in Ukraine, is mainly a secondary cavity nester. After egg-laying, chicks start to fly within 26 to 28 days, but still depend on their parents for about 3 weeks more. The diet of adult rollers is dominated by *Coleoptera*, whereas nestlings mostly eat *Orthoptera*, such as grasshoppers and bush crickets (Catry et al., 2019); opportunistically small vertebrates and smaller insects (e.g. winged ants) are also consumed (Sosnowski, Chmielewski, 1996; Aviles, Parejo, 2002; Kiss et al., 2014). Autumn migration begins in August/September.

The European range of the roller was formerly more extensive, but there has been a long-term decline in numbers and in range (Birdlife International, 2015), particularly towards the north of the range, e.g. Poland (Sosnowski, Chmielewski 1996) and Estonia (Lüütsepp et al. 2011), and much of NW Ukraine (Havrys’, 2009). These marked population declines have been accompanied by local extinctions and overall range contraction due to, as suggested, land use changes (Kovács et al. 2008) and/or anthropogenic climate change causing historically unprecedented rates of transformation (Huntley et al., 2006), or, more likely, the interplay between both these gross factors. On the ground the underlying cause of recent and ongoing declines of species is the growth of human populations and associated impacts ranging from habitat loss up to physiological stress responses to variable levels of human activities (Expósito-Granados et al., 2020), nest predation and killing or taking of birds (Havrys’, 2009; Brochet et al., 2016; Belyalova, 2020; etc.). With an overall decline of more than 30% the roller was classified as ‘vulnerable’ in Europe (Burfield, van Bommel 2004), although a later assessment reclassified the species as of ‘least concern’ (Birdlife International, 2015). In the Red Data Book of Ukraine (Havrys’, 2009) the species is listed under the category “Declining”.

According to the ‘International Species Action Plan for the European Roller’ (Kovacs et al. 2008), threats identified as ‘critical’ to the European roller are: intensification of forest and grassland management, increased habitat homogeneity, conversion of permanent grasslands, land abandonment and increased insecticide use. Amongst other potential threats climate change has been named. Yet there are large gaps in the study of these threats (Finch, 2016). In summary, they have led to both food and nest-site limitation, suggested as being responsible for the decline of the European roller (Kovacs et al. 2008). Consequently, a key part of many conservation interventions is the provisioning of artificial nest-boxes (Rodriguez et al., 2011), a management measure largely missing in Ukraine. The installation of nest-boxes for rollers proved to be an efficient method to replace the lack or the loss of natural nesting sites (Avilés et al., 2000), while other studies reported lesser success of nest-box programs (Sosnowski, Chmielewski, 1996), indicating most likely the insufficience of food sources, namely large-bodied insects. A recent study showed that, approximately 10 years before extinction of the local population in southern Poland, European rollers preyed on large beetles and orthopterans, which is typical, but recent agriculture abandonment and natural succession may have reduced the abundance of these prey species and their availability for the birds, thus contributing to their extirpation in the area (Hebda et al., 2019). Over the last few decades, orthopterans, for example, have experienced a considerable decline in Europe, with practices such as insecticide application, intensive cutting and grazing, and the spread of rationalized grass monoculture having detrimental effects (Wilson et al., 1999). In another study employing a species distribution model to explore the effects of potential climate change scenarios on the distribution of the wart-biter bush cricket (*Decticus verrucivorus*), an important food item in the diet of the roller (Hebda et al., 2019), a prediction was made that under severe climate change, the cricket will be left with very little suitable habitat in Europe (Carne, 2017). This highlights the threat of climate change to the species and suggests the same may apply to other species in the roller’s diet.

In general, recent findings using species distribution models hypothesize that future climate changes will likely amplify the impacts of existing threats on the majority of large roller populations in Europe (Kiss et al., 2020). In our study, we too used a species distribution modeling approach (Austin 2002, Guisan, Thuiller 2005) to hindcast changes in the climatically suitable range of the roller during historical times across Ukraine and to derive spatially explicit predictions of climatic suitability for the species under current climate.

Species distribution models, or SDMs, closely related to ecological niche models, bioclimate-envelope modelling etc., generally correlate species’ occurrence patterns with environmental variables. Especially SDMs have shown to be efficient in biodiversity research considering climate changes (Barbet-Massin et al., 2011; Visconti et al., 2016). As input, SDMs require georeferenced biodiversity observations. Often these occurrence points vary in geography and show significant gaps in the literature, even within one country. Therefore a prerequisite for our study was to collate all published and unpublished data to build a comprehensive dataset of occurrences of breeding pairs of the roller in Ukraine.

The specific goals of this study were to reconstruct the historical climatically suitable range of the species in Ukraine before the 1980s, gain better knowledge on its climatic requirements, compare the past and current climatically suitable areas, infer the regional and environmental variables that best explain its occurrence, and quantify the overall range change (‘loss’ or ‘gain’) in the country.

## Materials & Methods

### Dataset

Building of the datase was based on bibliographic research, checking curated repositories of specimens in Ukrainian museums and institutions, mailed questionnaire surveys, consultations with fellow ornithologists, and on personal field work. To map the retrospective distribution of breeding rollers, we analyzed the collections of 5 leading museums in the country. In this case, only those specimens of the collections were considered that were obtained during the breeding period. The collection of the Zoological Museum of the National Museum of Natural History (NASU) (Peklo, 1997) contains 42 specimens of the roller, caught in 17 regions of Ukraine; the collection of the Zoological Museum of the Taras Shevchenko National University of Kyiv – 21 specimens from 7 regions; the collection of the State Museum of Natural History (NASU) in Lviv (Bokotey, Sokolov 2000) – 12 individuals from 5 regions of the western region of Ukraine; the collection of the Zoological Museum of Uzhgorod National University (Potish, Potish 2006) – 1 specimen caught in the Transcarpathian region; the collection of the Museum of Nature of the V.N. Karazin Kharkiv National University – 25 individuals from 8 regions of the country (information kindly presented by T.M. Devyatko). All specimens of the collections were caught in the period from 1851 to 1972. In addition to the collections, materials of the Department of Zoology of the Taras Shevchenko National University of Kyiv were processed, kindly provided by Prof. V.V. Serebryakov: information from 83 questionnaires from 24 regions of Ukraine for the period of 1960–1986 were used. Questionnaire data of the Ukrainian Society for the Protection of Birds, collected in the framework of the awareness action “European Roller – Bird of the Year (1995)”, was kindly provided by the secretary of the society T. Mikhalevich: 49 questionnaires were received, our analysis used data from 44 questionnaires coming from 16 regions of the country. In terms of bibliography, 37 literary sources on the distribution of the roller in Ukraine were analyzed. In this respect recently published 3 volumes of “Materials for the 4th edition of the Red Book of Ukraine” (2018a,b; 2019) are an indispensable source of updated information concerning years 2010–2020. As part of a dissertation work the “Ecology of the Birds of the orders *Coraciiformes* and *Upupiformes* in the Ukrainian Steppe)” (Shupova, 2010) occurences of breeding pairs, using conventional methods (Novikov, 1953), were collected in the field between 1991–1997.

Geographical coordinates were determined using the location details provided in the label, or in the literary source, and employing the GeoCalculator in the DIVA-GIS program (Hijmans et al., 2001).

### Environmental Data

A severe challenge for modeling the temporal changes concerning suitable ranges of the European roller in Ukraine based on a lengthy record of occurrences extending back to 1851 is the inconsistency or absence of sets of available environmental variables in a GIS format that reflect historical parameters for certain time periods (Cohen et al., 2019). Sometimes the available data is limited in space or of unexceptably coarse resolution. These shortcomings significantly hamper modeling efforts and the search for factors responsible for the decline of roller populations.

Identifying the key environmental variables that determine the niche is one of the most crucial in SDM operations. Organisms usually respond to a complex of interdependent factors that consist of many environmental variables (Rydgren et al. 2003). Long ago Joseph Grinnell (1917) listed the factors that potentially affect the species distribution, e.g. vegetation, food, climate, soil, breeding and refuge sites, interspecific effects, and species preferences. However, it is often difficult or impossible to find more or less complete sets of those variables, particularly of biotic character. Fortunately, remote sensing and geographical information system (GIS) technologies provide a wide spectrum of spatial information that assist in the evaluation of macro-distribution of species (such as climate, land-cover etc.) (Martínez Pastur et al., 2016).

Firstly, in this study we used factors such as temperature and precipitation, since they are related to processes and impacts that are central to the persistence of the bird species. Importantly, also they can be assumed to be central to the well-being of the bird’s prey.

TerraClimate is a global gridded dataset of meteorological and water balance variables for 1958-present, available on a monthly timestep (Abatzoglou et al., 2017). Its relatively fine spatial resolution, global extent, and long length are a unique combination that fills a void in climate data. These data can be used in species distribution modeling, to approximate local variability and changes where station-based data are lacking or derived variables are preferred, and for climate-impact analyses in ecological systems cases where spatial attributes of climate may be preferred over coarser resolution data. The raw TerraClimate netcdf files are available for download by clicking on the file link https://climate.northwestknowledge.net/TERRACLIMATE/index_directDownloads.php.

Using SAGA GIS (Conrad et al., 2015), spatial resolution of the original rasters was resampled to ~1 km, as this satisfyingly approximates the home range sizes of nesting rollers (Finch, 2016; Finch et al., 2019).

We utilized a set of factors that were hypothesized to be of importance to roller presence and securing a sufficient food base. 1) Abiotic factors, such as temperature (tmin, tmax) and precipitation (ppt), were employed because they are consistently found to be primary determinants of species distributions at broad scales (Wiens, 2011). 2) Potential evapotranspiration (pet) can be especially informative for understanding broad-scale ecological patterns (Fisher et al., 2011), created by the synergy of temperature, humidity, solar radiation, wind, and biomass (Baltensperger, Joly, 2019). 3) Actual evapotranspiration (aet) was used in order to assess the dependency on plant resource availability, which was previously described as enabling successful predictions of bird species richness, by replacing the Normalized Difference Vegetation Index (NDVI) with aet measures as the two are significantly correlated (Cohen et al., 2019). Moreover, recent findings support the applicability of NDVI data as a suitable habitat-specific proxy for the food availability of insectivores during spring (Fernández-Tizón et al., 2020). For instance, foliage cover is important for egg laying in the orthopteran *Calliptamus italicus*, one of the most frequent diet items of the roller in Ukraine (unpublished data), as eggs are laid in open areas with bare ground between vegetation patches (Uvarov, 1977). Conversely, medium-to highgrowing herbaceous layer provides a good windbreak and reduces the risk of the eggs dehydrating in the soil (Poniatowski, Fartmann, 2008). 4) Downward shortwave radiation (srad) is a primary energy source in ecosystems (Yang et al., 2010), significantly affecting land surface processes (e.g., ecological, hydrological, biogeochemical) (Liang et al., 2010), which interacting with soil moisture (soil) and climatic water deficit (def) shape vegetation patterns (Bonan, 1989). In addition, soil properties such as soil moisture have been shown to influence oviposition, and therefore further offspring viability in grasshoppers (Herrmann et al., 2010). Also solar radiation can affect the performance of insect herbivores (Battisti et al., 2013) and their reproductivity (Bale et al., 2002). In grasshoppers, for instance, solar radiation is an important factor for body temperature regulation (Pepper, Hastings, 1952). 5) Finally, from the TerraClimate dataset we extracted wind speed (ws). Wind is a key climatic variable for flying birds (Cornioley et al., 2016). It potentially affects a wide range of activities from foraging to migration (Shepard et al., 2013). In particular, wind influences foraging efficiency of birds by modulating energy expenditure and movement speed (Hedenstrom, Alerstam 1995). Together with solar radiation, wind speed defines specific microclimates and their effects on water and energy budgets of birds and are of major importance to our understanding of avian thermal biology (Wolf, Walsberg, 1996).). Wind speed can also regulate local ambient temperatures, enhance the loss of soil moisture, facilitate long-distance dispersal of insects, representing potential prey of the roller (Yadav et al., 2018).

Besides meteorological and water balance conditions, changes in land cover and land use, including density of human population, significantly affect ecosystem processes including the carbon cycle, the water cycle, species diversity, and socioeconomic development. Here too, inconsistent time series representation is a problem. Fortunately, in some way the Global Land Analysis and Discovery (GLAD) Laboratory in the Department of Geographical Sciences at the University of Maryland (https://glad.umd.edu/) has filled this gap. Firstly, by producing a dataset representing a globally consistent cropland extent time-series (Potapov et al., 2021). Cropland expansion is known to have severe adverse effects on natural biodiversity (Pimm, Raven, 2000) through loss and fragmentation of habitats (Foley et al., 2005). The crop mapping was performed in five-year intervals (2000-2003, 2004-2007, 2008-2011, 2012-2015, and 2016-2019), however for our purpose the net cropland extent change from 2003 to 2019 was considered; pixel values (0-100) represent the percent of cropland dynamic (net loss or net gain) per pixel. Secondly, an annual vegetation continuous field product was developed consisting of tree canopy (TC) cover, short vegetation (SV) cover and bare ground (BG) cover, characterizing land change over the past 35 years (1982-2016) (Song et al., 2018). Three global map layers represent net changes in TC, SV and BG, respectively. Pixel values (−100 to 100) represent net percent change over the 35-year period. Negative values represent loss; positive values represent gain; zero represents no change.

Further, raster layers were employed for characterizing human population density known to impact biodiversity (Luck, 2007). In a static case we used the ‘Gridded Population of the World’, Version 4 (GPWv4) (Center …, 2018) for the year 2000, whereas for the study of human population density dynamics a high resolution global gridded data set representing a time series from 1981 to 2014 was employed (Lloyd et al., 2017).

In the end, we considered the velocity of climate change, a surprisingly elegant analytical concept that can be used to evaluate the exposure of organisms to climate change (Loarie et al., 2009; Garcia et al., 2014; Hamann et al., 2015). The measure is derived by dividing the rate of projected climate change in units of °C per year by the rate of spatial climate variability. The resulting variable is a speed or velocity measured in units of km/year, and represents an initial rate at which species must migrate over the surface of the earth to maintain constant climate conditions. In this study we performed the climate-analog velocity algorithm developed by Hamann et al. (2015) that calculates both distance and speed of the climatic parameter from present to the future climate match, using the aggregated data from the TerraClimate database for the time periods 1961-1990 and 1981-2010.

### Modelling

There exists a large suite of algorithms for modelling the distribution of species (Li, Wang, 2013; Hallgren et al., 2016). To explore the distribution of the European roller in our study area we employed Bayesian Additive Regression Trees (BART), a machine learning technique consisting of a Bayesian approach to Classification and Regression Trees (CART), capable of producing highly accurate predictions without overfitting to noise or to particular cases in the data. Models of this method estimate the probability of a given output variable (a binary classification of habitat suitability or species presence) based on decision “trees” that split predictor variables with nested, binary rule-sets (Carlson, 2020). Running SDMs with BARTs has recently been greatly facilitated by the development of an R package, “embarcadero”. The algorithm computes habitat suitability values ranging from 0, for fully nonsuitable habitat, to 1, for fully suitable habitat. It includes an automated variable selection procedure being highly effective at identifying informative subsets of predictors. Also the package includes methods for generating and plotting partial dependence curves, illustrating the effect of selected variables on habitat suitability. These response curves consist of the specific environmental variable as the x-axis and, on the y-axis, the predicted probability of suitable conditions as defined by the output. Upward trends for variables indicate a positive relationship; downward movements represent a negative relationship (Baldwin, 2009).

In terms of discrimination accuracy model performance was evaluated using two commonly used validation indices: the area under a receiver operating characteristic (ROC) curve, abbreviated as AUC, and the True Skill Statistic (TSS). The AUC validation statistic is a commonly used threshold independent accuracy index that ranges from 0.5 (not different from a randomly selected predictive distribution) to 1 (with perfect predictive ability). Models having AUC values >0.9 are considered to have very good, >0.8 good and >0.7 useful discrimination abilities. The TSS statistic ranges from −1 to +1 and tests the agreement between the expected and observed distribution, and whether that outcome would be predicted under chance alone. A TSS value of +1 is considered perfect agreement between the observed and expected distributions, whereas a value <0 defines a model which has a predictive performance no better than random (Allouche et al. 2006). TSS has been shown to produce the most accurate predictions (Jiménez-Valverde et al., 2011). Values of TSS < 0.2 can be considered as poor, 0.2–0.6 as fair to moderate and >0.6 as good.

Because of variable sampling intensity, occurrence points required by SDMs often vary in spatial density. As a result, and to avoid overemphasizing heavily on sampled areas, the BART algorithm selects points for model calibration using subsampling to reduce sampling bias and spatial autocorrelation, which would produce models of lower rather than higher quality (Beck et al., 2013).

We used the 10^th^ percentile training presence threshold value to generate binary maps (Liu et al., 2005). This threshold value provides a better ecologically significant result when compared with more restricted threshold values (Phillips, Dudík, 2008) or more liberal ones. Based on the probability value, we divided the study area into 2 classes: unsuitable and/or marginal area below the threshold value, in other words, where the species has predominantly gone extinct (also known as extirpation area), and an area where suitability is above the threshold value (chiefly habitation area).

Maps of habitat suitability in the GeoTIFF format were processed and visualized in SAGA GIS, statistical data was analyzed using the PAST software package (Hammer et al., 2001) and/or the R environment (R Core Team, 2020). If necessary, raw data was log-transformed or the Box-Cox transformation was applied.

## Results

### Presence records

The update of published and unpublished data yielded a total of 347 of non-duplicate records georeferenced occurrences: 148 for the period prior to 1980 (Fig. 1), 51 for records made between 1985 and 2009 (Fig. 2), and 148 records made between 2010 and 2021 (Fig. 3).

**Fig. 1.**
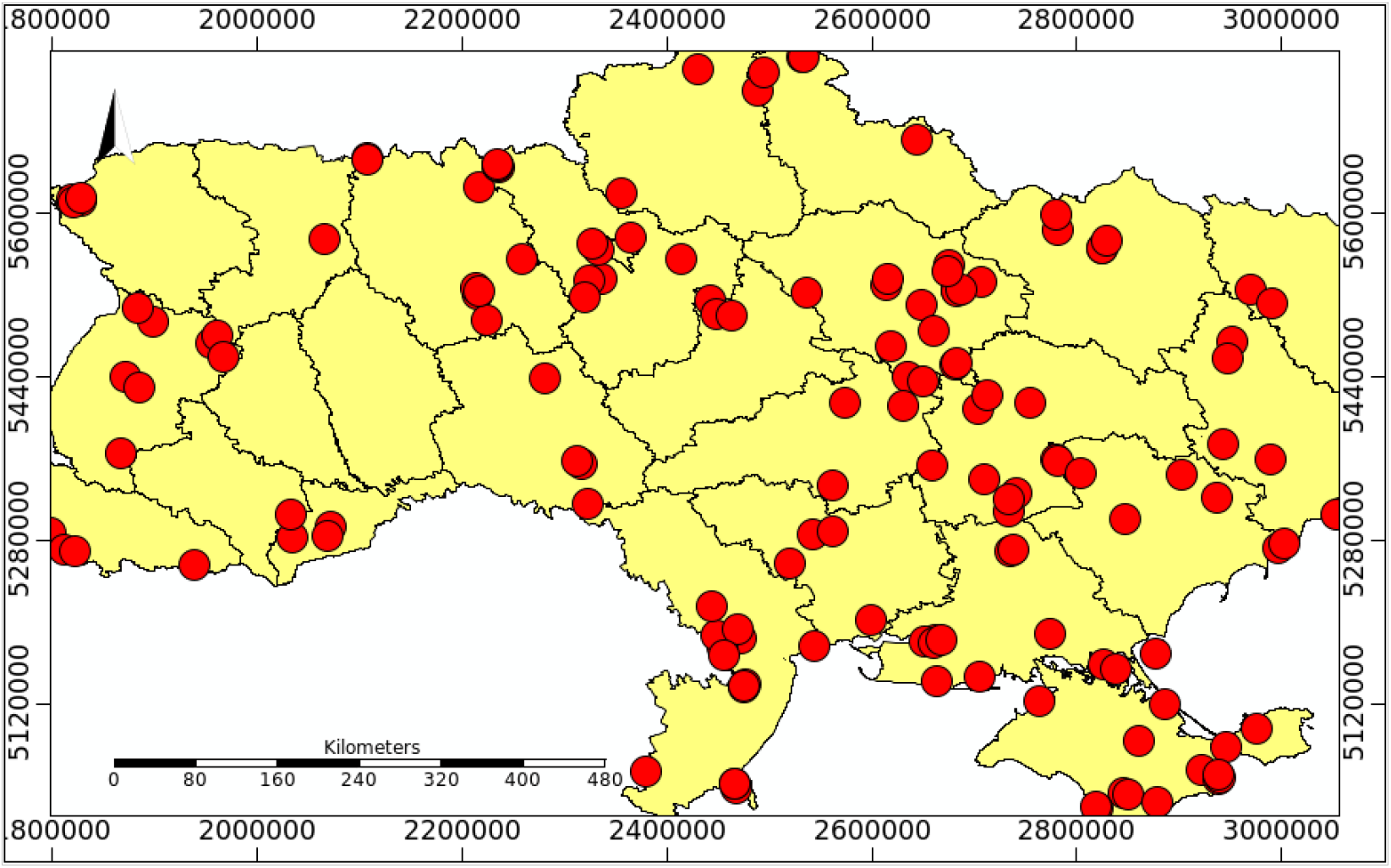
Distribution of the European roller in Ukraine before 1980 (red dots)

**Fig. 2.**
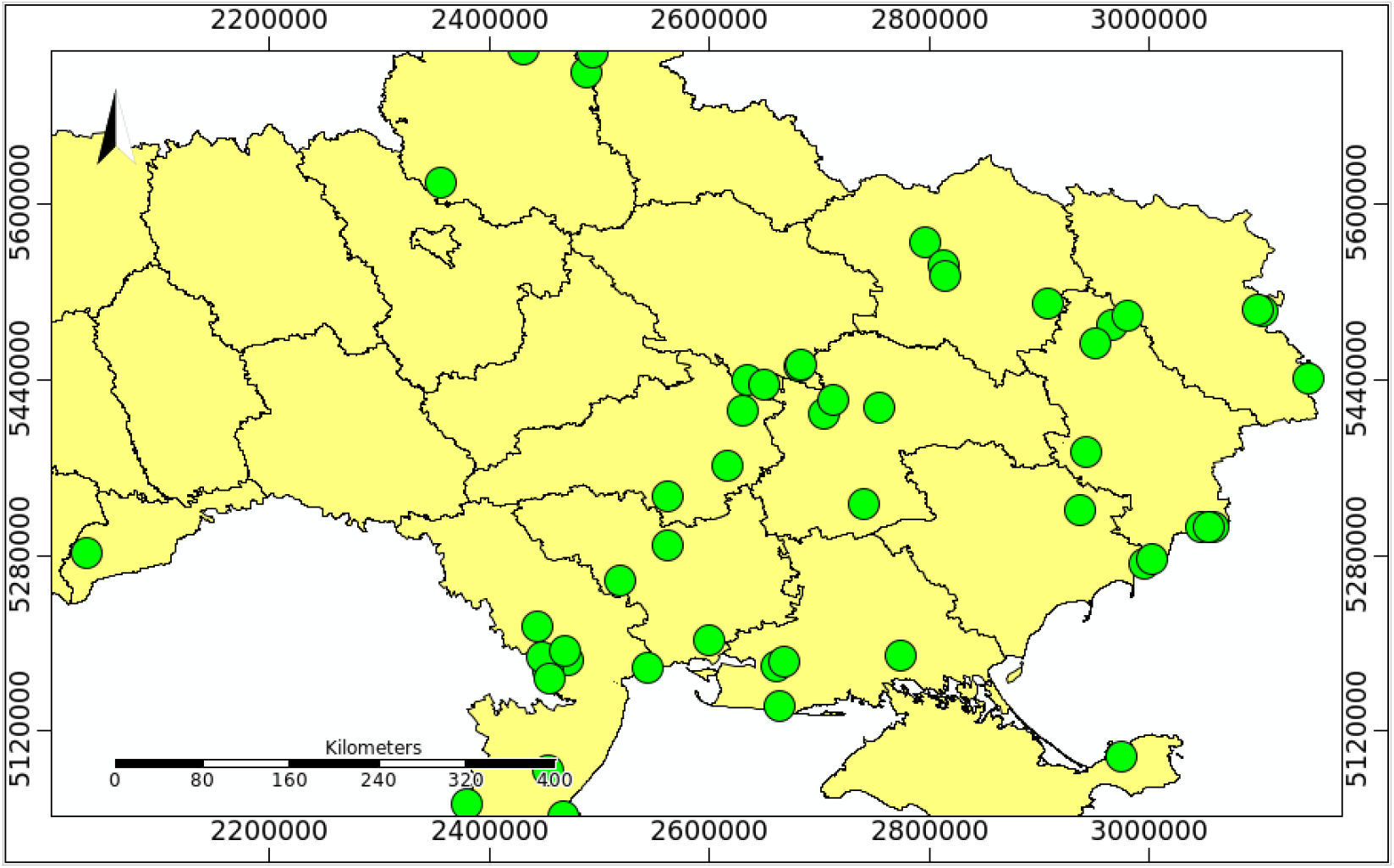
Distribution of the European roller in Ukraine between 1985 and 2009 (green dots)

**Fig. 3.**
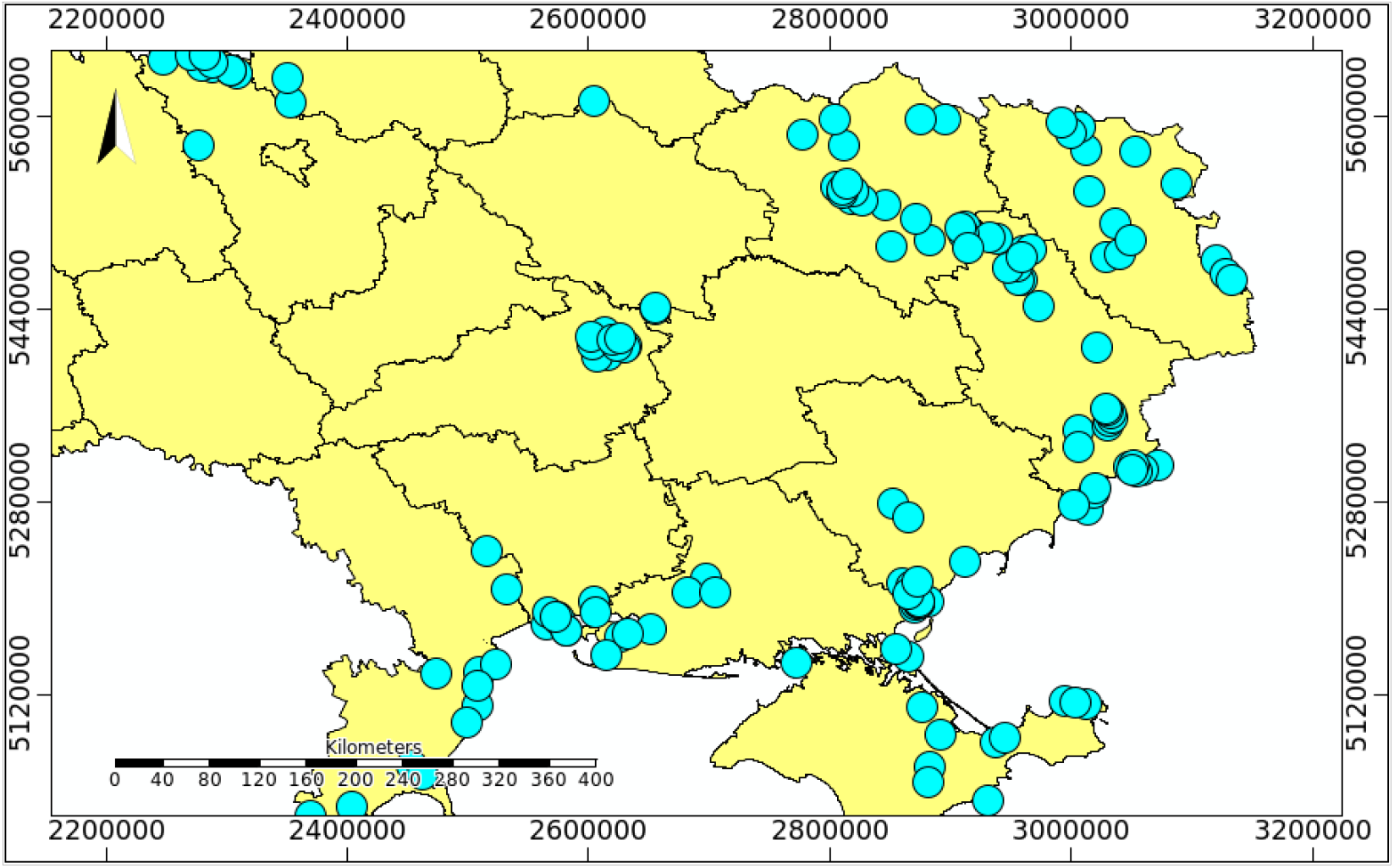
Distribution of the European roller in Ukraine between 2010 and 2021 (blue dots)

The shifting of the boundaries of the roller’s range to the east and south obviously became a trend in the 1980s. By 2021, a clear picture appears of the birds having localized in the eastern (east of Kharkiv, Luhansk, Donetsk) and southern regions of Ukraine, and regions adjacent to the Black and Azov seas, and in Eastern Crimea. In Donetsk and Luhansk regions, even in the 80s, when the decline in the number of the species was happening everywhere, the roller was not a rare bird.

A subpopulation of the roller, which until 1980 nested around Kyiv, by 2021 shifted to the area of the Chornobyl exclusion zone. In the 1980s and 1990s the roller had practically disappeared from the Kyiv region (Peheta, 1991). This is confirmed by the data in Fig. 2. For this period, a single record of the species was noted on the border of the Kyiv and Chernihiv regions.

### BART model performance

SDMs were created for the three time intervals (before 1980, between 1985-2009, and 2010-2021) using the corresponding climate data extracted from the TerraClim database for both the breeding season (May-August) and year-round conditions.

All BART models did a good job of differentiating known localities from background points (0.87 ≤ AUC ≤ 0.95; 0.59 ≤TSS≤ 0.78). Similarity between habitat suitability models accounting for breeding season and year-round conditions was high (the coefficient of determination, *R^2^*, varying from 78.82% to 88.78%; p<0.05), however based on expert knowledge we considered the breeding season models to give a better representation of both the past and current distribution of the European roller in Ukraine, and moreover, in neighbouring areas (see below). Correspondingly, three habitat suitability maps were produced: for the period prior to 1980 (AUC = 0.86, TSS = 0.59)(Fig. 4), years between 1985-2009 (AUC = 0.95, TSS = 0.78), and the years between 2010-2021 (AUC = 0.93, TSS = 0.73). Because there was a close similarity between the 1985-2009 and 2010-2021 models (*R^2^* = 75.60%; p<0.05), we decided to pool the data and build a joint habitat suitability model for this whole time period (AUC = 0.93, TSS = 0.73) (Fig. 5).

**Fig. 4.**
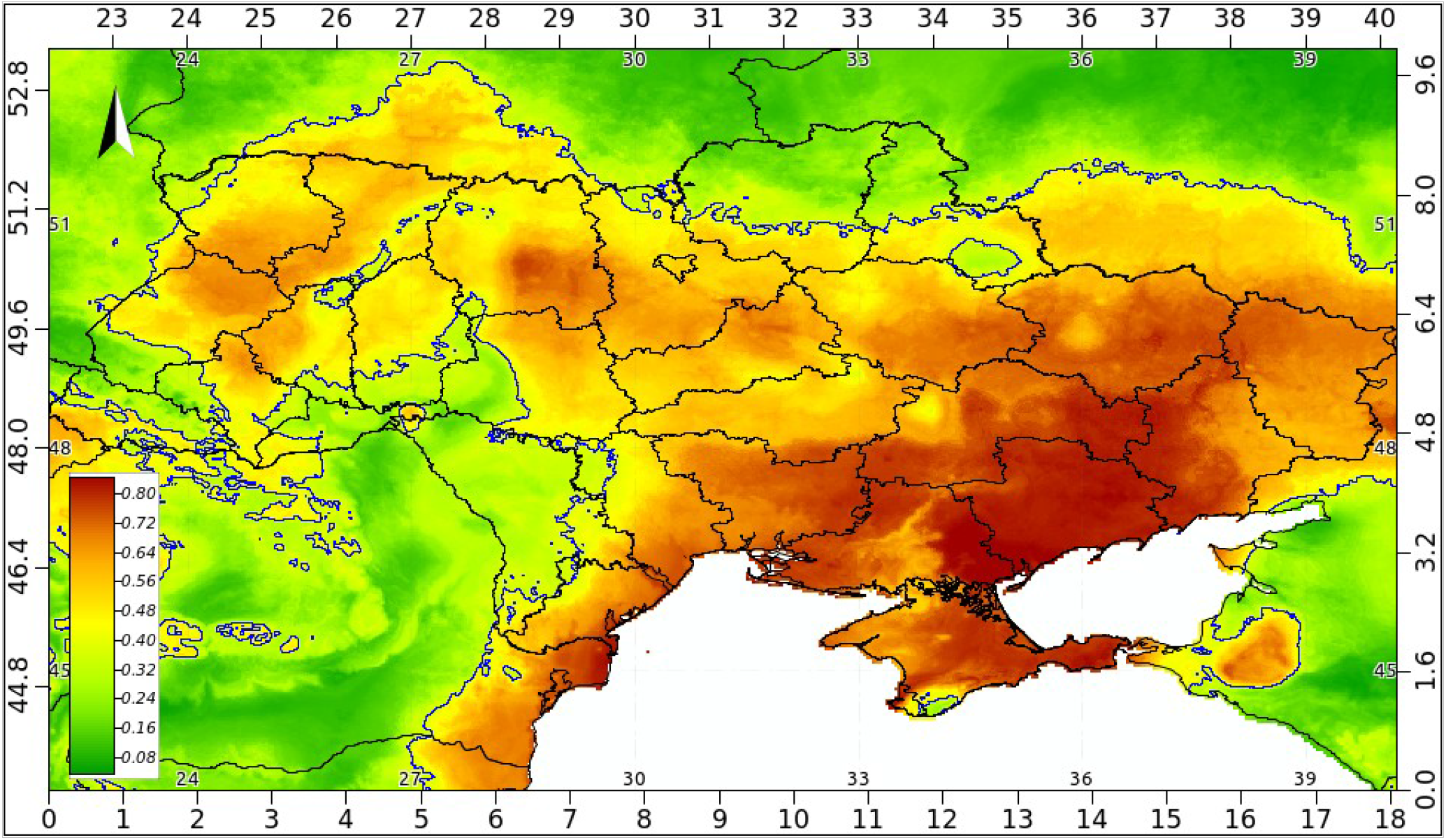
Habitat suitability map for the European roller in Ukraine built for the period prior to 1980; the legend shows habitat suitability ranging from high (brown) to low (green).

**Fig. 5.**
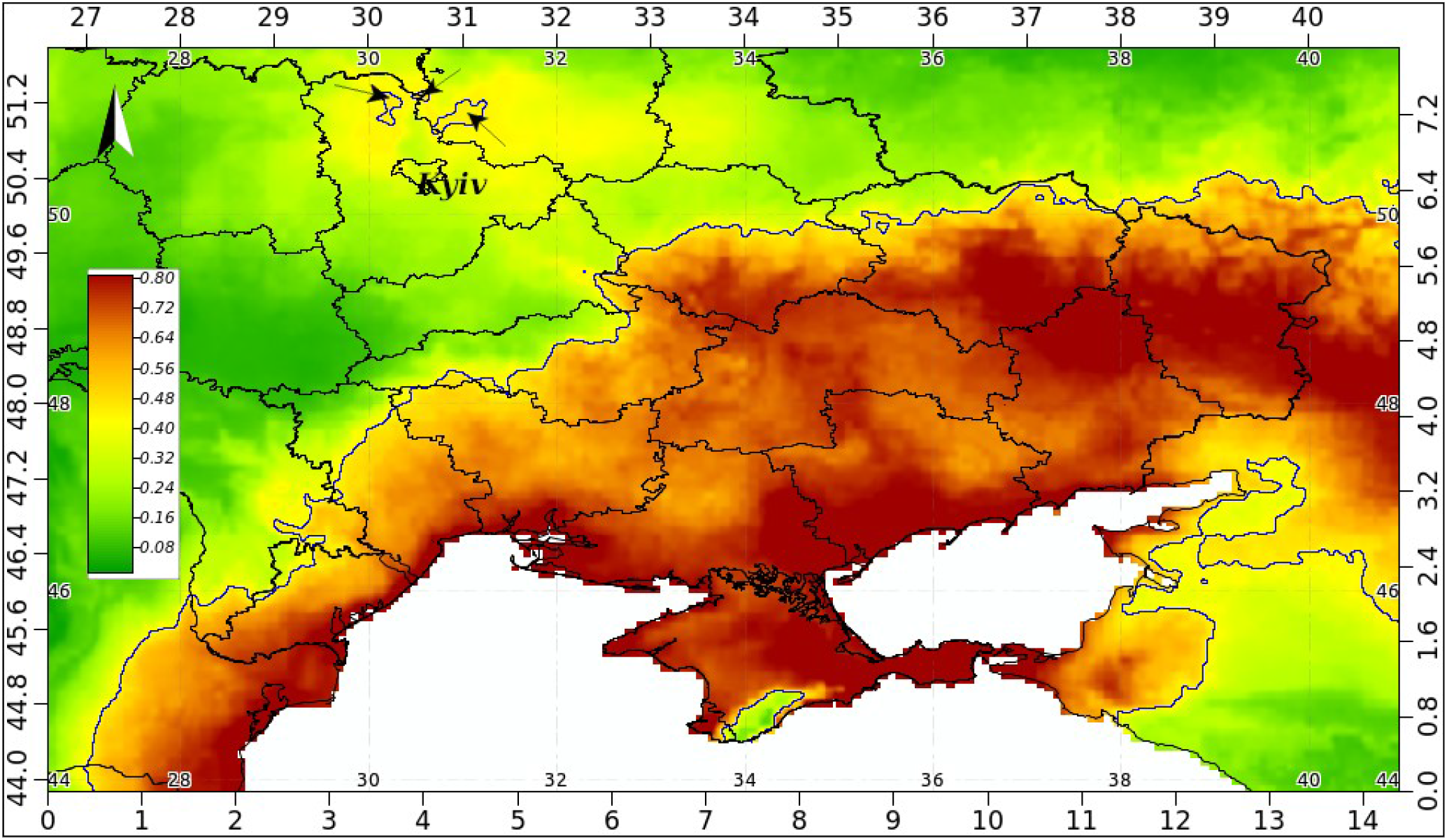
Habitat suitability map for the European roller in Ukraine built for the period 1985-2021; arrows point to enclaves north of Kyiv (including the Chornobyl Exclusion Zone); legend as in Fig. 4.

The obtained SDMs were reclassified as unsuitable areas (0–40%), low potential habitat suitability (40–50%), medium potential habitat suitability (50–70%), and high potential habitat suitability (70–100%). We defined these thresholds based on Martínez Pastur et al. (2016) proposals. In total, potentially suitable for the roller areas in Ukraine before the decline of the species comprised around 85%, whereas after the decline it was reduced to 46%. Areas with a low and moderate potential suffered noticeable losses, 25 and 34.7%, respectively. However, there was a gain in areas with a high potential for accommodating the roller – up to 60.2%, primarily in the east and south of the country, including Crimea (but with the exception of the majority of the southern coast and adjoining mountains). Despite this gain, the overall average habitat suitability in the country has fallen from 0.553 to 0.415 (Student’s t = 90.3; p<0.05).

The four most significant variables affecting the distribution of the roller in its Ukrainian range before the decline were in order of decreasing magnitude, average downward shortwave radiation (‘srad’), average actual evapotranspiration (‘aet’), average soil moisture (‘soilm’), and average maximum temperture (‘tmax’); partial dependence plots (i.e., response curves) for each of these are depicted in figs. 6 – 9. Other variables, like wind speed (‘ws’), were dropped from the model by the BART algorithm as insignificant. Not very surprisingly, the same set of variables are responsible for forming the bioclimatic niche of the roller in its contemporary area of habitation in Ukraine and their corresponding partial dependence plots appear to be of similar shape. From these response curves conclusions can be made that breeding rollers prefer a fairly narrow spectrum of the heat influx, areas of short and/or sparse vegetation, dry soil, average breeding season maximum temperature reaching 21.6°C, after which habitat suitability shows a steep drop (Fig. 9).

**Fig. 6.**
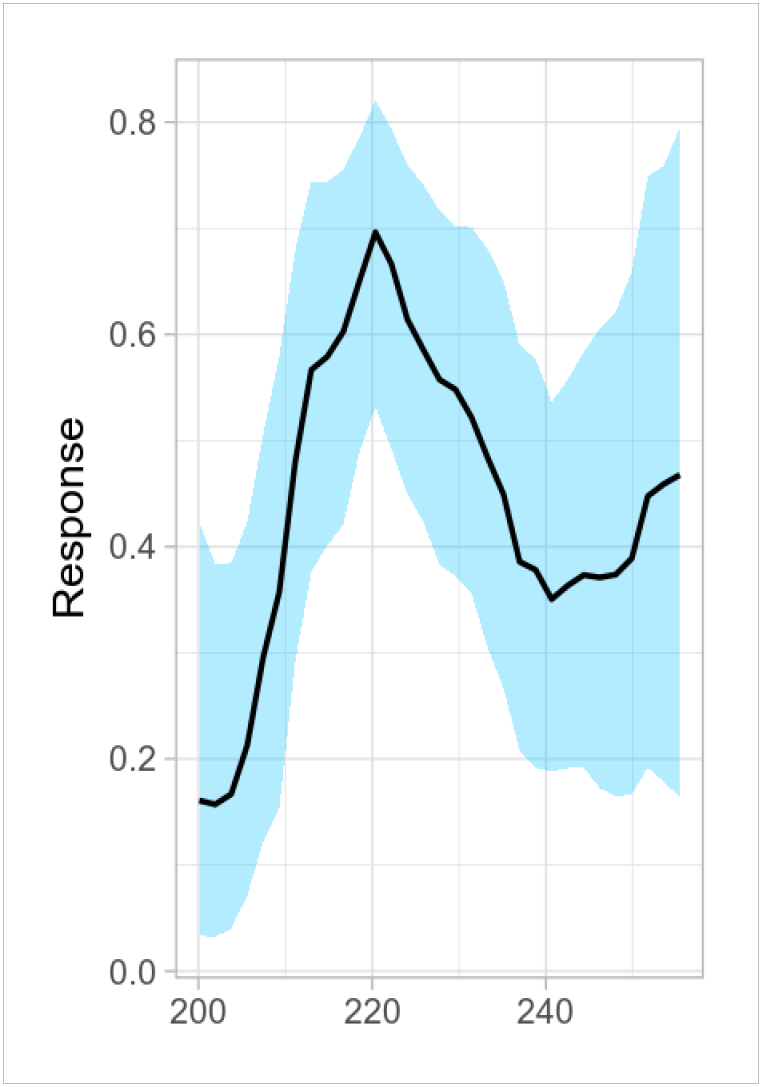
Partial dependence plot for average downward shortwave radiation (‘srad’); units = W/m^2^. Response = habitat suitability score, blue area = 95% confidence interval.

**Fig. 7.**
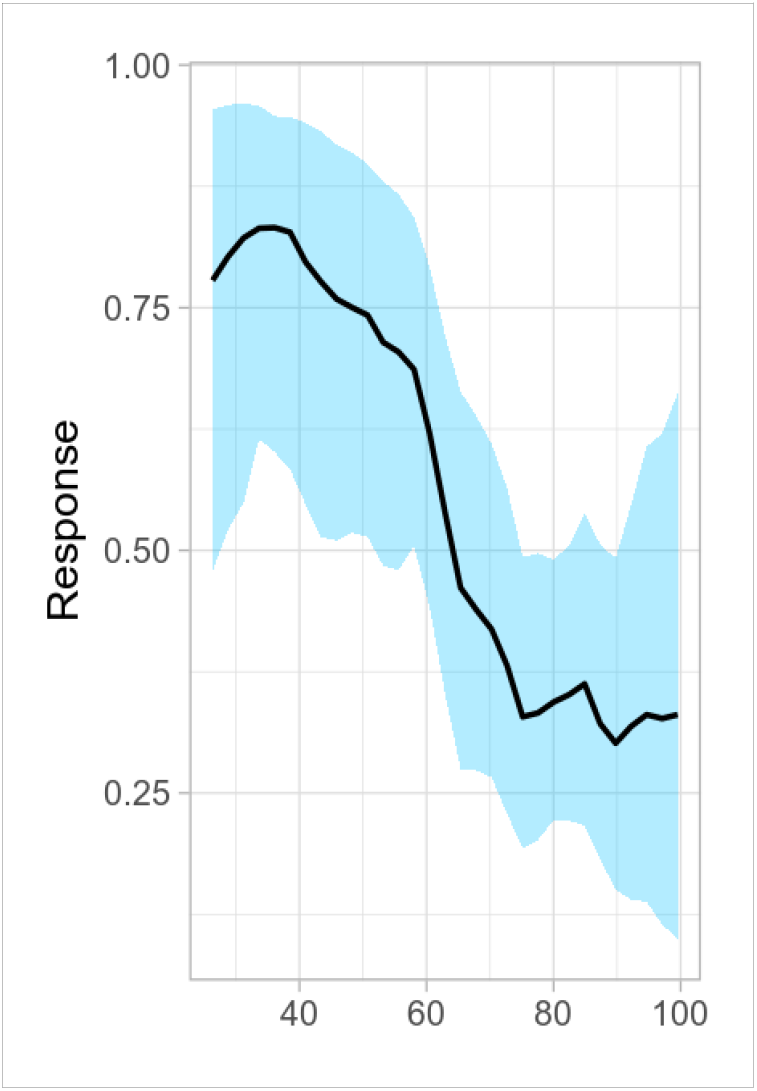
Partial dependence plot for average actual evapotranspiration (‘aet’); units = mm.

**Fig. 8.**
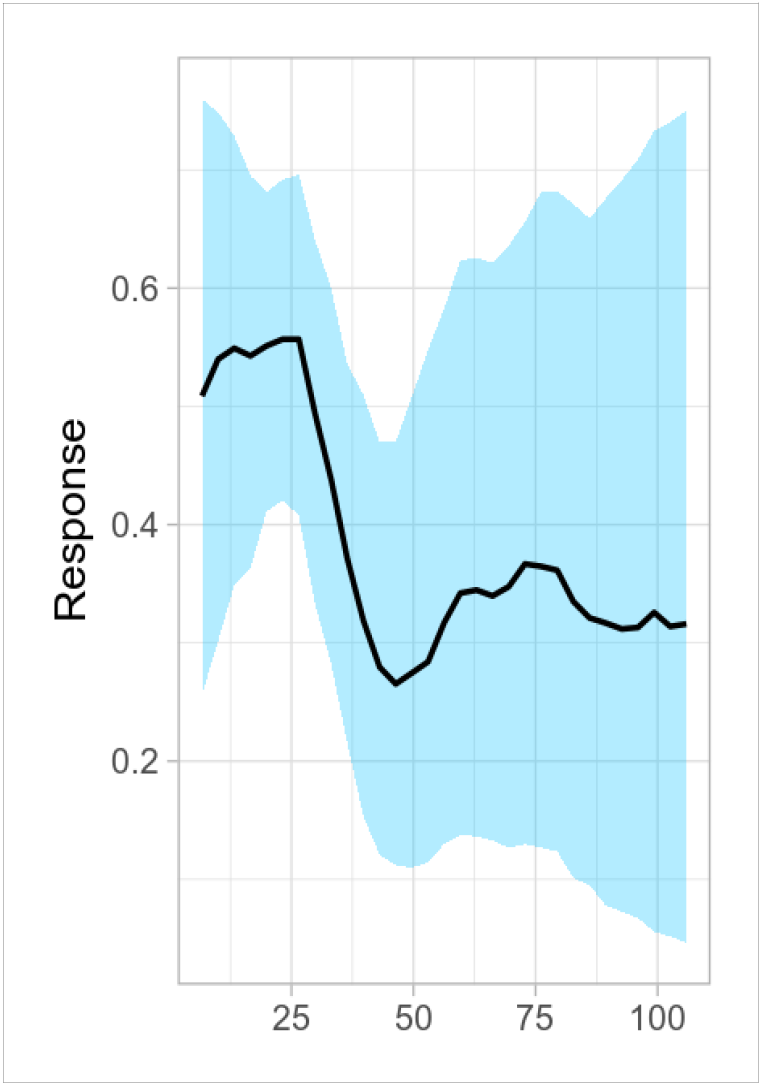
Partial dependence plot for average soil moisture (‘soilm’); unit = m^3^/m^3^.

**Fig. 9.**
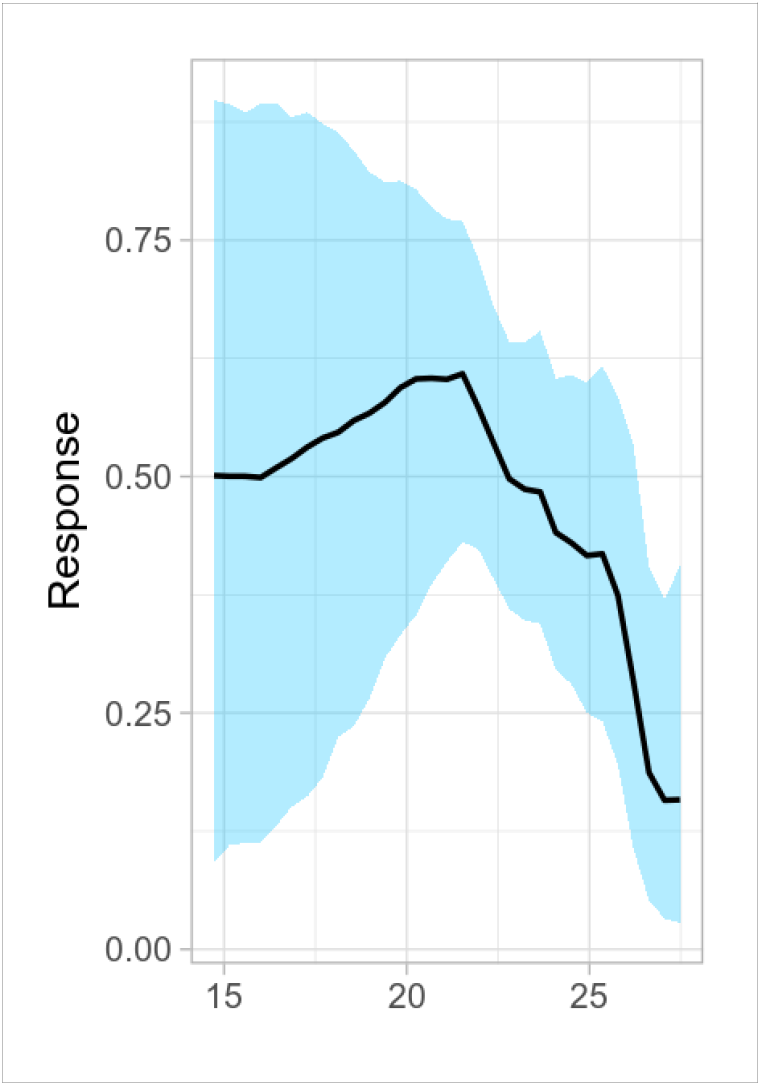
Partial dependence plot for average maximum temperature (‘tmax’); unit = °C.

### Changes in land cover and land use

Using the 10^th^ percentile threshold, we made a distinction between the extirpation area (predominantly in the NW of the country) and the area of contemporary habitation (mainly in the SE), and compared land cover and land use dynamics, both within their boundaries and/or between them, focusing on the surroundings of roller breeding sites as they were prior to 1980, after when the decline of the roller in the study area began.

### Cropland dynamics

Both within the extirpation area and area of contemporary habitation and between them cropland dynamics seem to have remained more or less constant and at a fairly low level (in the range from 4.1 to 5.6%), with loss and gain apparently compensating one another.

### Vegetation field dynamics

Dynamics of the annual vegetation continuous field products mentioned above are presented in Table 1.

**Table 1.** Dynamics (1985-2016) of annual vegetation continuous field products in areas of extirpation of the roller (since 1980) and its contemporary habitation.

| Vegetation field | Extirpation area | Habitation area | Student's t (p<0.05) |
| --- | --- | --- | --- |
| Bare ground | -0.78 | -1.59 | 2.6 |
| Short vegetation | -10.52 | -0.50 | 7.7 |
| Tree canopy | +11.99 | +2.64 | 6.9 |

Looking at the ‘bare ground’ category, one can see losses have occurred in both considered areas. More striking changes have affected the other two categories, showing a noticeable reduction of ‘short vegetation’ and significant gain of ‘tree canopy’ cover in the area today mainly abandoned by breeding rollers.

### Human density

According to the census of the year 2000, human population density in the surroundings of nesting rollers was higher in the contemporary habitation area of the species, 114.4 against 101.8 ind. per sq. km in the extirpation area (Student’s t = 2.2; p<0.05). In terms of the dynamics of human population densities recorded from 1981 to 2014, no growth has taken place within the breeding sites of the roller from which the bird today is absent (Student’s t = 1.1; p>0.05), whereas in the contemporary habitation area of the species human population densities have increased (regression slope 7.13±1.13, Student’s t = 2.7; p<0.05).

### Velocity of climate change

Various climate surfaces can be used to generate estimates of the velocity of climate change, and include various biologically relevant temperature and precipitation variables, as well as extremes, growing and chilling degree days, various dryness and indices and growing season descriptors such as frost-free days (Hamann et al., 2015). We used the average maximum temperature for the breeding season, a seemingly crucial factor in the biology of the roller (see above). Once again, focusing on breeding sites as they were prior to 1980, we found a faster change of this factor in the extirpation area than in the area inhabited by the species today, 0.140 against 0.106, respectively (Student’s t =3.2; p<0.05).

## Discussion and conclusions

There is significant evidence showing that birds, as other animals (Parmesan, Yohe, 2003), are shifting their ranges in response to climate change. Both range expansions and contractions are occurring worldwide, although some bird species may remain unaffected by climate change, however range contractions are expected to be more frequent than range expansions (Wormworth, Mallon, 2006). Here the European roller is just one example of such range contraction presumably due to climate change. Between 1970 and 1990, the roller was declining in a majority of European countries (Tucker et al., 1994) and this pattern continued throughout the next decade. However, in 2000–2010 the roller’s decline looked to have decelerated (Finch, 2016). Our modelling exercises seemingly support this view: a comparison of the model created using occurrences recorded prior to 1980 with the model for the 1985–2009 time period shows low similarity (R^2^ = 46.4%), whereas similarity between the latter and model for the years 2010–2021 appears significantly higher (R^2^ = 75.7%), meaning fewer changes have taken place within this time period regarding the distribution and parameters of bioclimatic habitat suitability of the species in Ukraine. Saying “fewer” does not, of course, mean that there were zero changes. Fortunately or not, they are continuing and our climatic predictions show that things can become worse.

In the meantime, the European roller in Ukraine has retreated to the south-west of the country and today its northern home range boundary (as defined by the 10 percentile threshold) closely follows the boundary of the Steppic biogeographical region (Cervellini et al., 2020) (Fig. 10), therefore characterizing the species in Ukraine as ‘steppic’ is largely justified.

**Fig. 10.**
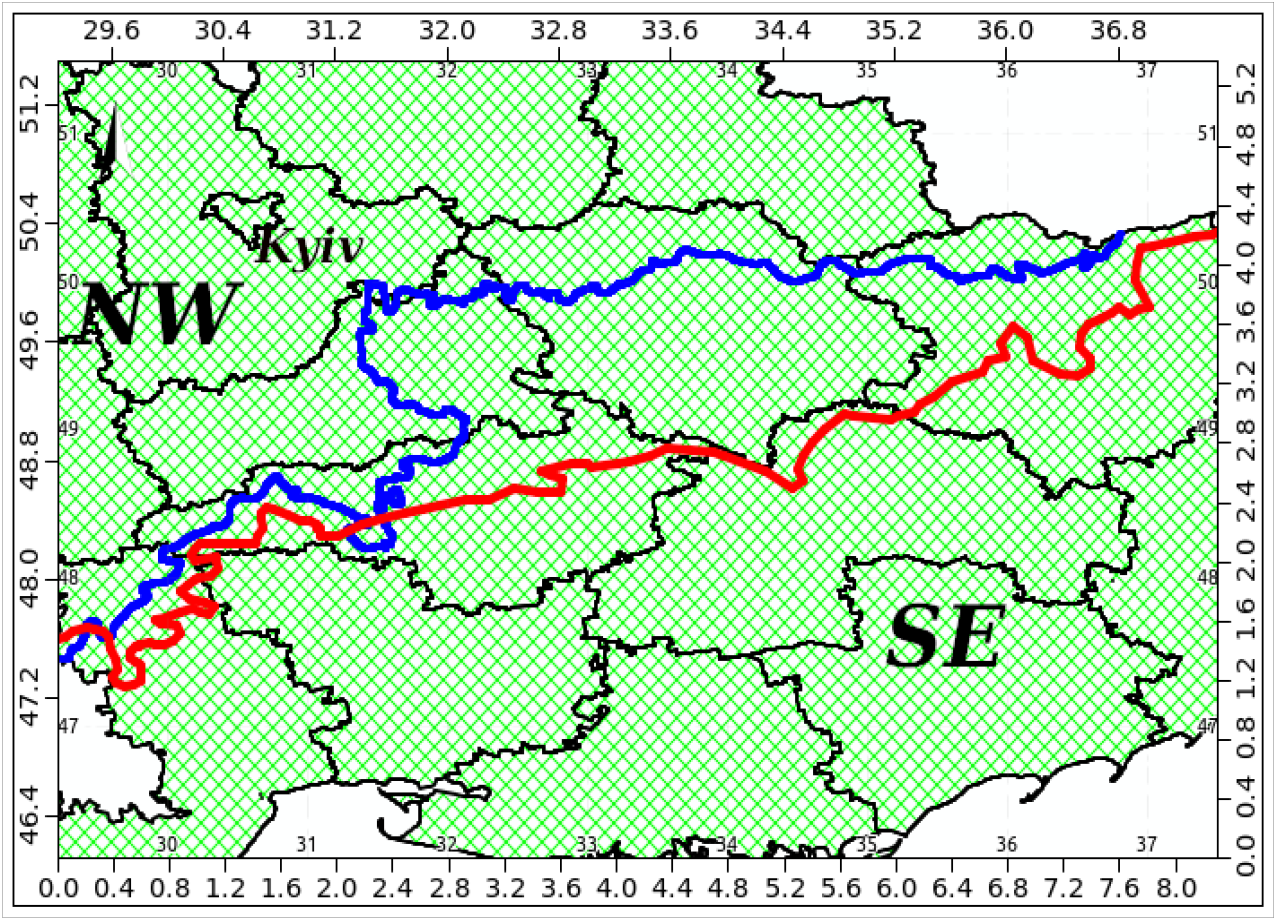
Spatial relationship between the 10 percentile threshold separating the extirpation area in the NW and contemporary habitation area of the European roller in the SE of Ukraine (blue line) and the boundary of the Steppic biogeographical region (red line).

As common practice, we assessed the predictive performance of our SDMs by randomly splitting the dataset into training and testing, and fitting the model on the training dataset and validating it on the testing dataset using the area under the curve (AUC) and the true skill statistic (TSS). In our case these two metrics revealed good results. However, discrimination accuracy metrics may suggest a very good model while of poor transferability (Torres et al., 2015) and/or lead to relationships without a biological meaning (Santini et al., 2021). Using only Ukrainian-based records, we tested our models by extrapolating predictions of occurrence and habitat suitability to neighbouring areas, for instance to Krasnodar Province of the Russian Federation (Fig. 11). These extrapolations showed close to excellent predictive power by pointing out sites most favourable for the roller, namely the Taman and Yeysk peninsulas, where within the region the bird is found in its greatest abundance (Red Data Book …, 2017). Response curves (Figs. 6–9) too, in our opinion support the credibility of our SDMs by producing results that have a biological interpretation consistent with the physiology and/or ecology of the bird species or either environmental requirements of its potential prey.

**Fig. 11.**
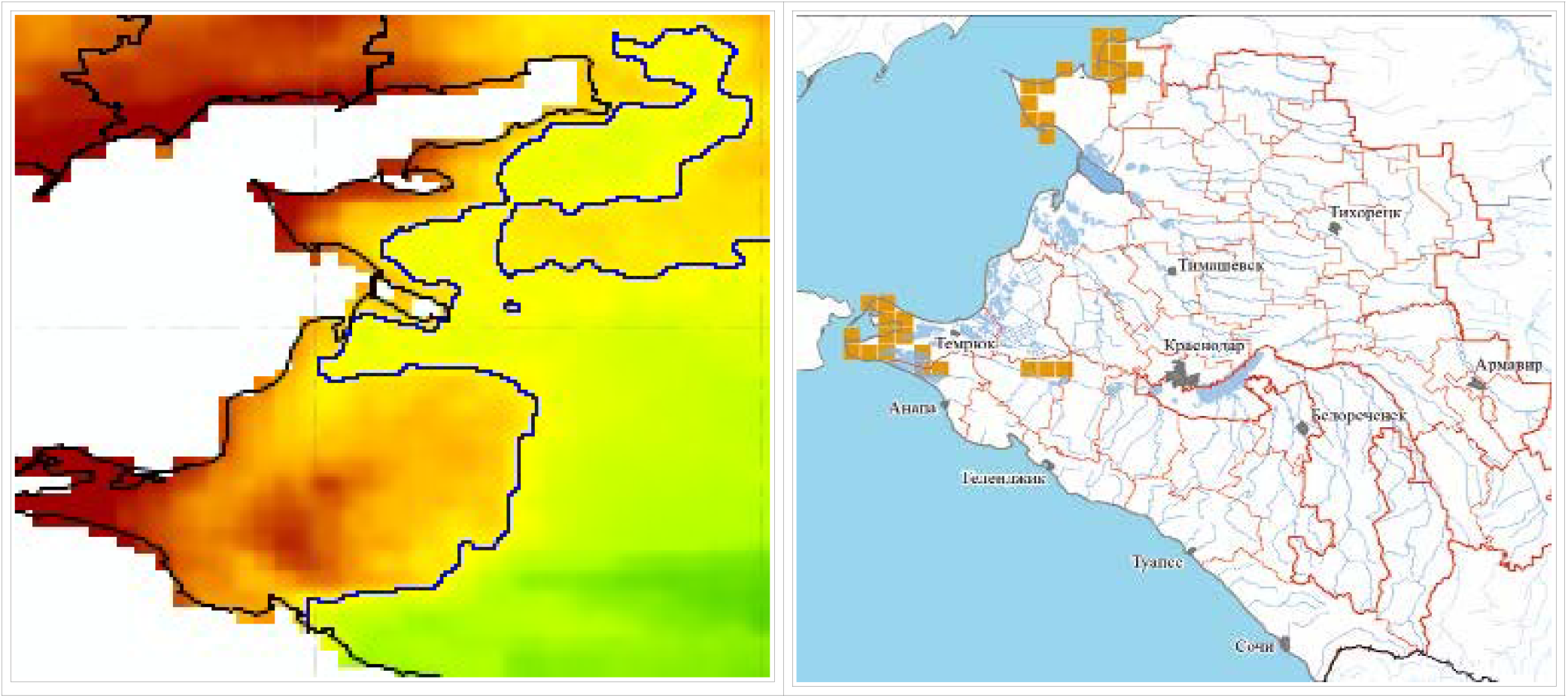
Left: extrapolation of the 1985-2021 species distribution model for the European roller to Krasnodar Province [habitat suitability ranging from high (brown) to low (green)]; right: map from the Red Data Book of Krasnodar Province (2017), p. 577, showing the distribution (squares) of the species in the region.

As mentioned above, one of the most crucial in SDM operations is identifying the key environmental variables that determine the niche of the species in question. Usually, SDMs are calibrated only with abiotic variables as predictors, assuming that biotic interactions are indirectly represented by abiotic variables because they strongly correlate (Soberón, Nakamura, 2009). The addition of biotic interactions usually improves the predictive performance of SDMs (Araújo, Luoto, 2007), however the inclusion of such interactions as, for example, the dependence of the roller on certain food items meets difficulties due to the insufficiency of appropriate data. Not able to incorporate abiotic and biotic predictors into one SDM, we considered building two: one for the roller and one for *Calliptamus italicus* (Linnaeus, 1758), the Italian locust, a species of ‘short-horned grasshoppers’ belonging to the family *Acrididae*, using the same set of climatic variables extracted from the TerraClimate database for the years 2010-2021 and employing the BART algorithm. Point data for the grasshopper was downloaded from the GBIF repository (GBIF.org, 2021). The choice of the prey species was determined by the fact that the Italian locust is a frequent food item for the roller in Ukraine: out of 172 individuals of *Acrididae* found in the stomachs of sampled birds 41 (or 23.8%) belonged to this species, more than it was for other grasshoppers (unpublished data). Assuming a coupling between the spatial distribution of resources and consumers, we hypothesized that the geographical distribution of the roller should in some way match that of the Italian locust and tested the expected spatial congruence across grid cells of the produced SDMs. To control for spatial non-independence we used a modified t-test to calculate the statistical significance of the correlation coefficient (a corrected Pearson’s correlation) based on geographically effective degrees of freedom as implemented in the SpatialPack package (Osorio, Vallejos, 2014). The relationship was found to be positive (0.487), statistically significant (p = 0.05) and can be considered moderate in its magnitude (Rumsey, 2016). Therefore, despite solely using abiotic variables as predictors, SDMs built for the roller roughly managed to capture features of the niche important for the locust too without explicitly introducing the insect to the model as a biotic factor. Perhaps, if the association between the bird and prey would be closer, let’s say the rollers would specialize and predominantly feed on locusts, the correlation coefficient would be higher. In the end, we found it reasonable to consider modeling the distribution only with abiotic variables and relying on the BART algorithm for identifying those to be of most importance.

Because of the gaps in the study of threats facing the European roller (Finch, 2016), we considered several factors that could have contributed to the decline of the species, namely in Ukraine, but maybe also in a broader geographical context.

Reasons for this decline are primarily focused upon the spread and intensification of agricultural systems leading to habitat fragmentation and loss (Saunders, 2016). Cropland (% of land area) in Ukraine was reported at 56.76% in 2018, according to the World Bank collection of development indicators (https://data.worldbank.org/indicator/), meaning much of the roller’s habitat in Ukraine is lost. Today minor changes in cropland dynamics within the country hardly have any impact on the welfare of the species.

An analysis of land cover trends more or less coinciding with the decline of the species in Ukraine bare ground losses, more pronouncedly in the SE of the country. Bare ground dynamics is an important component of global land cover change resulting from economic drivers such as urbanization and resource extraction (Ying et al., 2017). Bare ground losses could mean more vegetated sites. The closure and/or reduced operations of industrial plants and agricultural processing facilities in the eastern part of Ukraine (Burakovsky, Betliy, 2009) seem to have facilitated this trend. Later it was shown that abandoned in the Donbas region quarrying and coal mining sites, with the encroachment of vegetation, significantly enhance biodiversty (Ulyura, Tytar, 2017, 2018).

Larger changes have been seen regarding the dynamics of ‘short vegetation’ and ‘tree canopy’ cover and these because of their magnitude are more likely to affect the roller. The loss of areas with short vegetation and increase in the tree canopy cover in the NW of the country, now primarily abandoned by the species, certainly will not favour its return. Perhaps an exception is the Chornobyl Exclusion Zone and some adjacent areas, where Landsat images show the change from a previously vibrant agricultural and forestry economy, when crops have been replaced by grasslands (https://www.usgs.gov/news/earthview-chernobyl-30-years-later).

As for human population density and dynamics, these unlikely can be responsible for the extirpation of the roller from the NW as far as the occupied by the species SE of the country is more heavily populated by humans and densities here between 1981 and 2014 have noticeably increased.

The majority of threats posed to the roller by habitat and land use change are also likely to be compounded by the effects of global climate change (Saunders, 2016) and in our study this is exactly what climate velocity points to. In fact, velocities are a simple function of spatial and temporal variation in climate conditions in a particular landscape, and can be interpreted as one of several risk factors that contribute to persistence or loss of species and populations in complex landscapes under climate change (Hamann et al., 2015). Ultimately, we suggest climate change and its speed have been responsible for shaping the contemporary home range of the European roller in Ukraine and likely in other countries of the continent.

